# Similarity of drug targets to human microbiome metaproteome promotes pharmacological promiscuity

**DOI:** 10.1101/2024.04.26.591258

**Authors:** Christopher A. Beaudoin, Shannon Norget, Sharif Hala, Andries J. van Tonder

## Abstract

Similarity between candidate drug targets and human proteins is commonly assessed to minimize the occurrence of side effects. Although numerous drugs have been found to disrupt the health of the human microbiome, no comprehensive comparison between established drug targets and the human microbiome metaproteome has yet been conducted. Therefore, herein, sequence and structure alignments between human and pathogen drug targets and representative human gut, oral, and vaginal microbiome metaproteomes were performed. Both human and pathogen drug targets were found to be similar in sequence, function, structure, and drug binding capacity to proteins in diverse pathogenic and non-pathogenic bacteria from all three microbiomes. The gut metaproteome was identified as particularly susceptible overall to off-target effects. Certain symptoms, such as infections and immune disorders, may be more common among drugs that non-selectively target host microbiota. These findings suggest that similarities between human microbiome metaproteomes and drug target candidates should be routinely checked.

## 1. Introduction

Drug discovery campaign success has improved throughout the past three decades[1]. In recent years, an average of approximately 50 drugs per year have been approved by the USA Federal Drug Administration Center for Drug Evaluation and Research for therapeutic use, which is an increase compared to the historic average of 34 drugs approved per year since 1993[2]. Nevertheless, developing novel and safe drugs poses several challenges and requires significant time and resources to validate clinical efficacy, toxicity, sensitivity, and specificity[3,4]. To expedite the process for discovering new drugs, several experimental and computational approaches have been developed to enable high throughput small molecule screening campaigns[5,6]. Furthermore, significant advances in genome sequencing technologies[7–9] and protein structure determination methodologies (e.g. X-ray crystallography[10], cryo-EM[11]) have provided profound insights into drug binding sites and inhibitory mechanisms for various human and pathogen proteins. In combination with *in vitro* and *in vivo* functional and phenotypic screens[12], these data have allowed for the generation of numerous sets of principles that may inform selection of protein targets for the rational design of therapeutic small molecules[13]. The rational selection of a new drug target for human diseases and microbial pathogens requires several considerations: binding pocket features (e.g. depth, electrostatics), binding site conservation, gene expression level, number of interacting pathways[14]. In all cases, the drug should be highly specific to one or specific targets in order to minimize off-target effects[15]. Checking the homology of the drug target to the human proteome is crucial to ensure that the functions of essential human proteins are not modulated[16]. Additional considerations in drug target selection may further enhance the success and safety of drug screening campaigns.

The human microbiome comprises all organisms living on the surface or within the human body[17]. The number of unique genes in the human microbiome has been estimated to be 100-150 times higher than the number of human genes[18,19]. The abundance and functional features of microbial partners have been reported to be important for the maintenance of health at specific body sites[20]. Microbiota are also sensitive to perturbations resulting from changes in diet, location, circadian rhythm, and numerous other endogenous and exogenous factors[21]. Notably, although the effects of antibiotics on the human microbiome have been extensively surveyed, numerous human (non-antibiotic) drugs have been discovered to both affect microbial health and be metabolized by the microbiome[22,23]. Additionally, the microbiome may metabolize drugs, thus changing the target-binding properties and off-target effects[19]. Although the composition of microbial species in a healthy microbiome differs between individuals from genetic and geographic backgrounds, gene function in the metagenomes have been shown to be largely consistent, suggesting that gene function may be more indicative of the roles that microbes play in host-microbe interactions[20]. Therefore, novel insights into drug target selection parameters that can be modulated to minimize off-target effects on the microbiome may provide safer treatment options.

Several studies have previously proposed methodologies for predicting the non-selective targeting of microbiota, e.g. using machine learning models that incorporate drug chemical properties and the microbial genome features[24,25]. However, to date, no comprehensive comparison of the sequence homology and structural features of the drug binding pockets for human and pathogen drug targets with proteins in the human microbiome has yet to be performed. Therefore, using sequence and structure based bioinformatics, the similarity and, thus, drug binding potential for 1,346 FDA-approved drugs that collectively target 739 human and pathogen drug targets[26] were assessed on representative metaproteomes of the human skin, oral, and gut microbiomes[27]. In summary, both human and pathogen drug targets were found to be highly similar in sequence, structure, function, and drug-binding capacity to microbiome drug targets. These results highlight the utility of checking sequence and structural homology of the phylogenetically diverse organisms in the microbiome to candidate drug targets for binding site selections. Further validation of microbiome drug target promiscuity may shed light on the etiology of unintended clinical outcomes.

## 2. Results

### 2.1. Sequence and functional similarities between drug targets and microbiome metaproteomes

Protein sequence homology has been widely used to infer structure and function of proteins. An amino acid sequence identity of over 30% has been suggested to be indicative of a common evolutionary ancestry and, thus, structure and a sequence identity of above 40-60% for shared function[28–32]. Protein sequence homology has also been used to infer promiscuity of ligand binding[33]. Recently, Santos *et al*. reviewed the mechanism of action of FDA-approved drugs and their corresponding protein targets[26]. Additionally, the MGnify database[27,34] has made available comprehensive datasets on the human gut, oral, and vaginal metagenome assembled proteomes. The protein sequences correspond to 289,026, 1,225, and 618 nonredundant metagenome assembled genomes (MAG) in the gut, oral, and vaginal microbiomes. Utilizing these combined data, the human drug target protein sequences were mapped to the gut, oral, and vaginal metaproteome sequences to find similarities in amino acid sequence and function. BlastP was used to perform global alignments between the drug target and metaproteome sequences[35]. Sequence similarity between drug target and microbiome sequences may reveal microbiome species that are most likely to be non-selectively targeted.

Among the total 737 drug target sequences, 126 total (77 human and 51 pathogen) were mapped with above a 30% global sequence identity to metaproteome sequences. Alignments of the microbiome metaproteomes and drug target sequences reveal differences in sequence identity distributions with respect to targeted organism (Figure 1A). The pathogen drug target sequences reported higher average sequence identity than those of the human targets to the metaproteome sequences in all three microbiome datasets. As expected, no human targets were found to be identical to any metaproteome sequences, while 174, 22, and 20 unique metaproteome sequences were found to be identical to pathogen targets in the gut, oral, and vaginal microbiome, respectively. Notably, the average sequence identity and number of identical matches between the pathogen drug targets and the gut metaproteome (average: 70.4%), in particular, were much higher than oral (48%) and vaginal (46.3%) metaproteomes. Sequence identity distributions comparing the human targets and the three metaproteomes were all similar.

**Figure 1.**
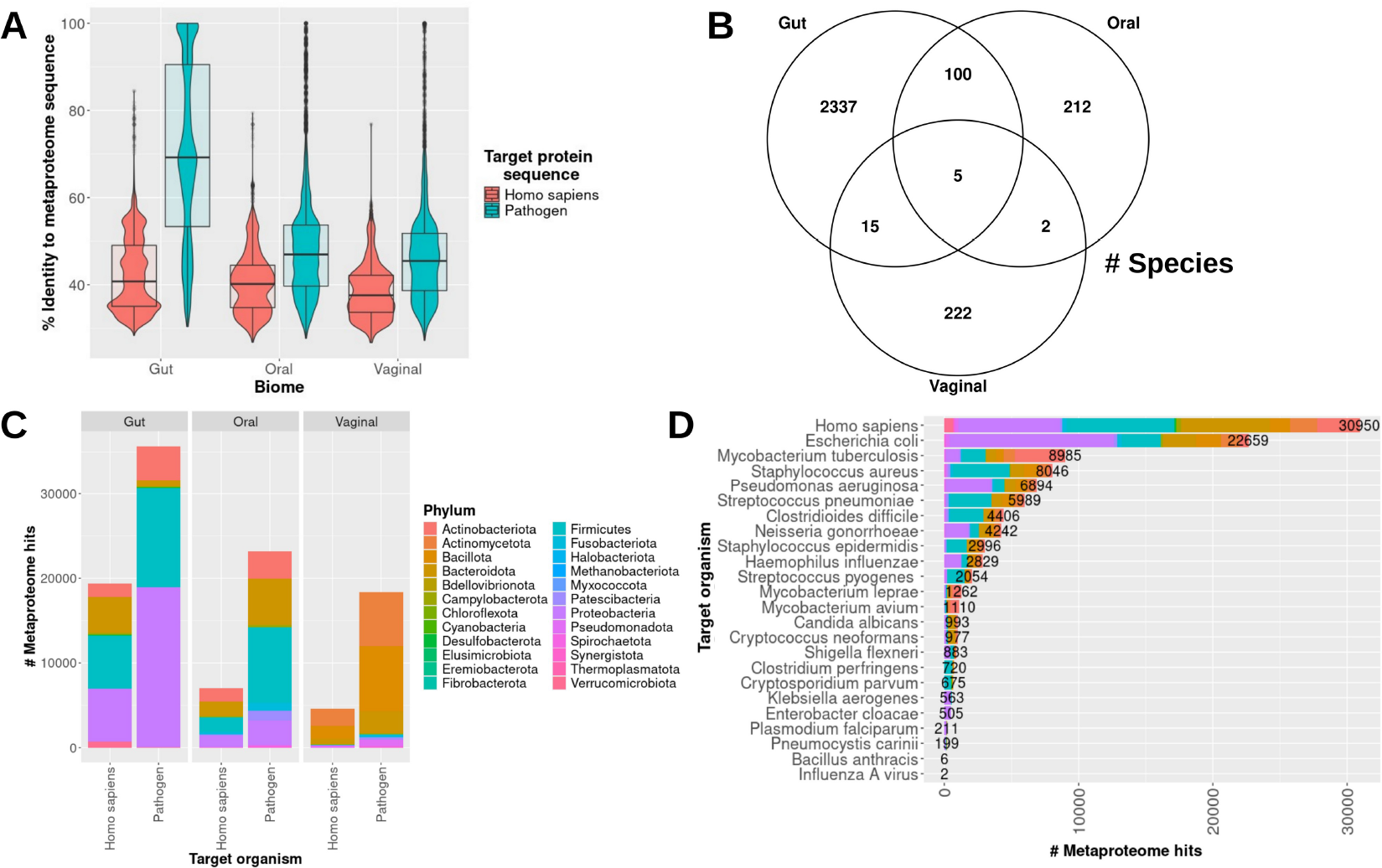
Sequence similarity between drug target and metaproteome sequences A combined violin and box plot showing the distribution of sequence identities between drug target and microbiome protein sequences for each metaproteome is shown (A). A Venn diagram highlighting the number of non-selectively targeted species shared between the microbiome data sets is portrayed (B). A bar plot showing the number of total similar genes between human and pathogen drug targets to the metaproteome sequences of each microbiome are shown and colored by microbiome protein lineage (C). Similar to (C), the numbers are shown per target organism (D).

More putative off-target species were found in common between the gut and oral metaproteomes than either of the two with the vaginal microbiome. Interestingly, 96% of the potential unintended targeted species were found to be specific to each microbiome, and only five species were found in common between all three microbiomes (Figure 1B). The primary affected phyla in the gut and oral microbiomes for both human and pathogen drug targets were Proteobacteria, Firmicutes, Bacteroidota, and Actinobacteriota, while the those in the vaginal microbiome largely comprised Bacteroidota, Bacillota, and Actinomycetota (Figure 1C). Although the sequence identity distribution profiles between the three metaproteomes and human drug target sequences were similar, the number of metaproteome sequences found to be potentially targeted in the gut microbiome (19,369) is much higher than the oral (6,980) and vaginal (4,601) microbiomes. The number of species affected by drugs for pathogen infections followed a similar trend to the human drugs across the three microbiomes: gut: 35,695, oral: 23,168, vaginal: 18,343. The highest number of off-target species was reported for the human targets (2,243), followed by those of *Escherichia coli* (1,146), *Mycobacterium tuberculosis* (1,060), *Clostridioides difficile* (990), and *Staphylococcus aureus* (863) (Figure 1D). In summary, phylogenetically diverse bacteria may be susceptible to drugs irrespective of the targeted organism.

Comparing the functions between mapped sequences may provide insights into which classes of drug targets might be differentially targeting metaproteomes. First, the functions of the drug targets were compared with the annotated functions of the metaproteome sequences, for which the data were available, to determine functional overlap. A sequence identity cutoff of over 50% was chosen based on previous findings, to reduce analytic constraints, and to validate sequence function comparisons[28,29]. Among human targets, alcohol dehydrogenase mapped only to S-(hydroxymethyl)-glutathione dehydrogenase, and the other metaproteome functional annotations all mapped to identical or similar functions (Figure 2A). The human target peptidyl-prolyl cis-trans isomerase protein sequence mapped to hypothetical proteins, peptidyl-prolyl cis-trans isomerases, and FK506-binding proteins in the metaproteomes. The annotated functions of the metaproteomes mapped to identical or similar functions for all pathogen targets (Figure 2B). The dihydrofolate reductase, 30S ribosomal protein S9, and 50S ribosomal protein L3 were similar in identity to hypothetical proteins; and dihydrofolate reductase mapped to IS1595 family transposases in the metaproteomes. Overall, the functional similarities between the mapped drug target and metaproteome sequences are evident. Therefore, an analysis of the species affected by drug target class may shed light on which drugs are most pharmacologically promiscuous.

**Figure 2.**
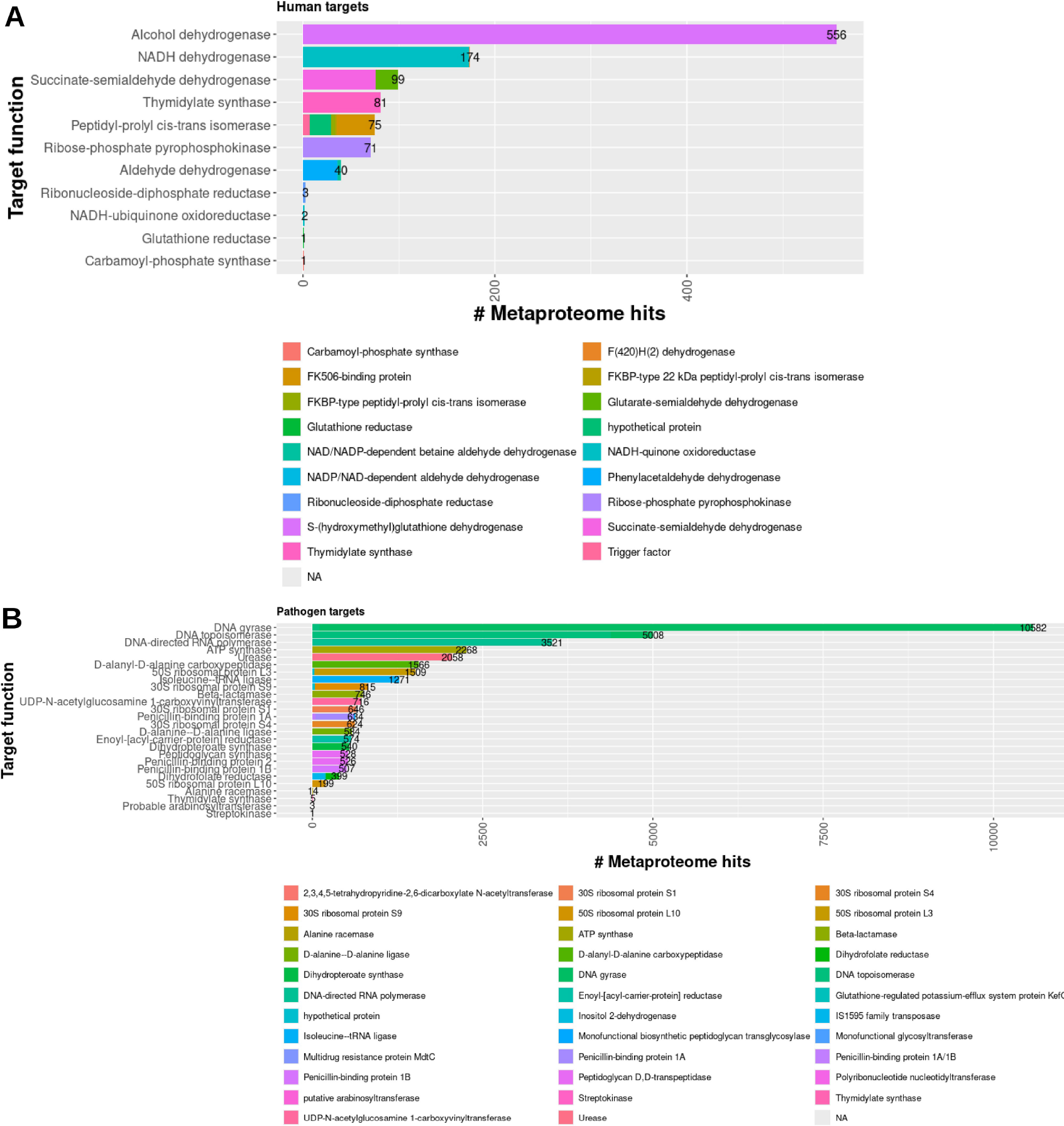
Relationship between target function and non-selectively targeted microbiome protein function Stacked bar plots representing the number of protein sequences were identified to be similar to human (A) and pathogen (B) drug target sequences are shown. The bars are colored by function of the microbiome proteins. The total number of protein sequences are shown at the end of each column to aid in the visualization of columns with smaller numbers.

The phyla non-specifically targeted by the human drugs are varied among the targeted proteins. As shown in Figures 3A and 3B, more phyla that represent the microbiome metaproteomes were found to be mapped to the human targets (24) than the pathogen targets (17). Most human and pathogen drug targets were found to be similar to Firmicutes, Proteobacteria, Bacteroidota, and Actinobacteriota, while five human drug targets (3-oxo-5-alpha-steroid 4 dehydrogenase, carbamoyl-phosphate synthase, neprilysin, dipeptidyl peptidase, sodium/glucose cotransporter 2) were highly if not exclusively similar to Bacteroidota. Proteobacteria are highly enriched in four pathogen drug target classes (D-alanyl-D-alanine carboxypeptidase, beta-lactamase, peptidoglycan synthase, and penicillin-binding proteins). The four pathogen drug targets (DNA gyrase, DNA topoisomerase, DNA-directed RNA polymerase, and ATP synthase) with the most similar metaproteome sequences were found to be nearly or more than double the number of all other human or pathogen genes, thus highlighting the increased potential for off-target effects. The sequence and function similarities between drug targets to the metaproteome proteins provide novel insights into the potential for drug promiscuity among the human microbiome. Additional investigation into the conservation of residues and structural features that are critical for drug binding may further clarify the capacity for off-target protein-ligand interactions.

**Figure 3.**
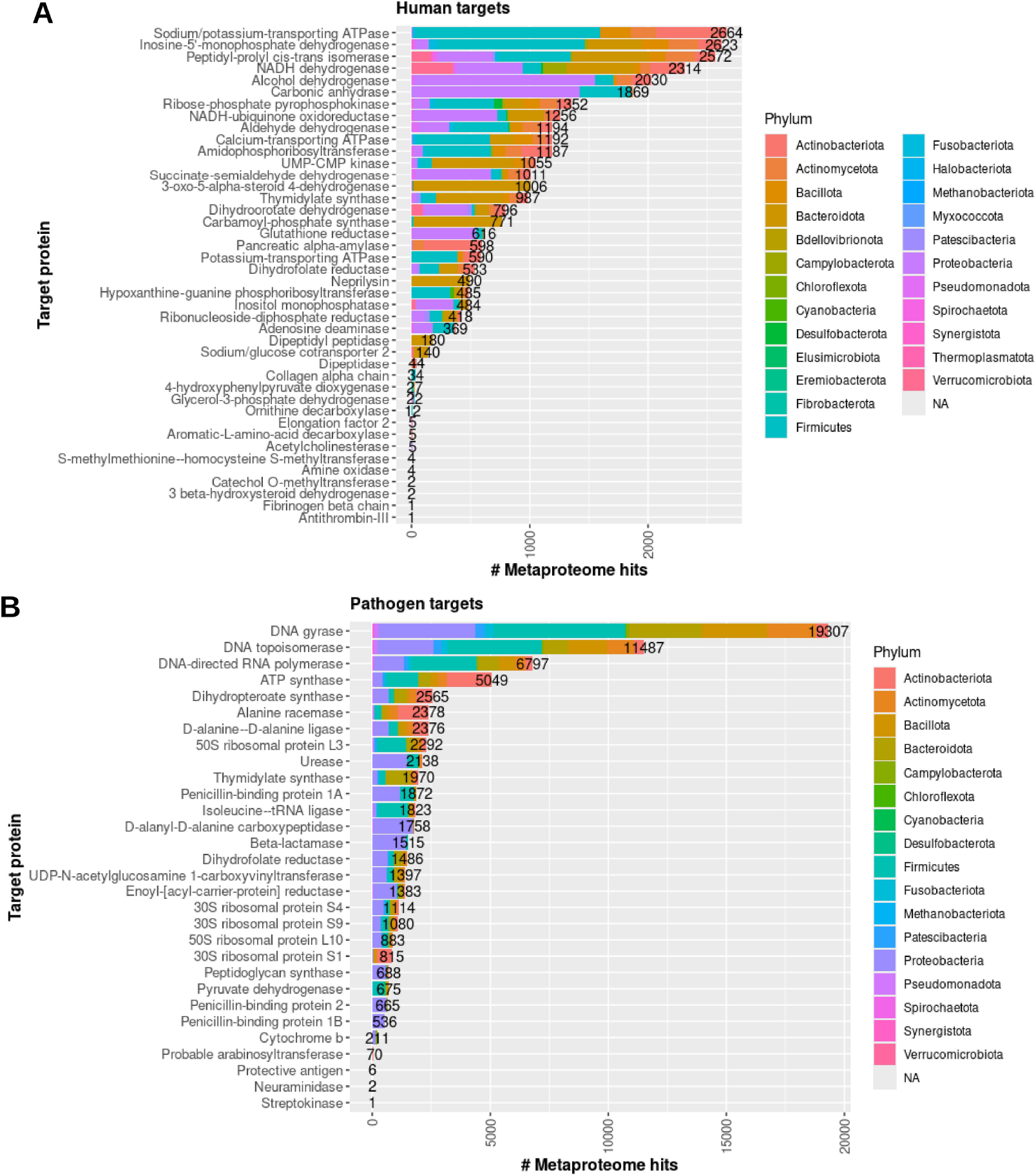
Relationship between target function and non-selectively targeted microbiome proteins Stacked bar plots representing the number of protein sequences were identified to be similar to human (A) and pathogen (B) drug target sequences are shown. The bars are colored by phylum of the MAG corresponding to the microbiome proteins. The total number of protein sequences are shown at the end of each column to aid in the visualization of columns with smaller numbers.

### 2.2. Structural and molecular similarities between drug targets and microbiome proteins

Protein-ligand interactions are largely determined based on the structural and electrostatic complementarity between the drug and binding pocket. Drug promiscuity has been attributed to similarity between drug binding pockets in different target proteins[36]. Therefore, examining the differences in drug binding pockets between the drug target and metaproteome protein structures may lend more insights into off-target interactions with the human microbiome. All experimentally-determined protein structures resolved with the exact small molecule from the drug target dataset were extracted from the RCSB Protein Databank (PDB)[37]. In total, four human and three pathogen drug-bound target structures were available for further analysis (Table 1). As shown in Figure 4A, the average sequence identity for the four human targets (41.8%) was lower than that of the three pathogen targets (54.0%). To practically test the drug promiscuity between phylogenetically distant organisms, the metaproteome sequence with the lowest identity to the drug target sequence was selected. Homology modelling of the microbiome protein structures was performed using the resolved drug target protein structures as templates. Subsequently, the corresponding drugs were docked into the metaproteome models based on the poses in the experimentally-determined drug target structures. The predicted affinity and protein-ligand interactions were generated and compared between the drugs with the microbiome and drug target proteins.

**Table 1.** Predicted protein-drug affinities for drug target and microbiome proteins.

|  |  |  |  |  |  |  |  |  | # Contacts |  |  |  |  |  |  |  |  |  |  |  |
| --- | --- | --- | --- | --- | --- | --- | --- | --- | --- | --- | --- | --- | --- | --- | --- | --- | --- | --- | --- | --- |
| Drug | Target organism | Protein_name | PDB | Target gene | Biome | % Identity | Species | $\Delta G$ (Kcal/mol) | CC | CO | CN | CX | OO | OX | NO | NN | NX | XX | | |
| Trimethoprim | *E. coli* | Dihydrofolate reductase | 7MYM | folA | - | - | - | -5.3 | 143 | 133 | 177 | 2 | 27 | 2 | 60 | 33 | 1 | 0 |  |  |
|  | - | - | - | - | Vaginal | 30.488 | Bifidobacterium vaginale | -5.51 | 138 | 134 | 177 | 3 | 26 | 4 | 59 | 27 | 3 | 0 |  |  |
| Ethambutol | *M. tuberculosis* | Probable arabinosyltransferase A | 7BVF | embA | - | - | - | -5.1 | 117 | 140 | 154 | 0 | 24 | 0 | 54 | 34 | 1 | 0 |  |  |
|  | - | - | - | - | Gut | 34.057 | Corynebacterium sp902373425 | -5.14 | 147 | 165 | 175 | 3 | 25 | 2 | 60 | 37 | 2 | 0 |  |  |
| Fidaxomicin | *C. difficile* | DNA-directed RNA polymerase | 7L7B | rpo | - | - | - | -15.58 | 4618 | 3086 | 1440 | 113 | 516 | 31 | 470 | 0 | 32 | 0 |  |  |
|  | - | - | - | - | Vaginal | - | Nanoperiomorbus sp004136275 | -15.94 | 4485 | 2986 | 1472 | 144 | 508 | 49 | 483 | 0 | 41 | 1 |  |  |
| Acetazolamide | *H. sapiens* | Carbonic anhydrase 2 | 6YMA | CA2 | - | - | - | -5.16 | 156 | 195 | 209 | 160 | 40 | 41 | 90 | 47 | 50 | 1 |  |  |
|  | - | - | - | - | Oral | 30.469 | Neisseria sp000952795 | -5.2 | 147 | 200 | 205 | 153 | 53 | 47 | 94 | 45 | 48 | 1 |  |  |
| Rapamycin | *H. sapiens* | Peptidyl-prolyl cis-trans isomerase | 6M4U | FKBP1A | - | - | - | -9.48 | 3135 | 1673 | 727 | 0 | 200 | 1 | 212 | 27 | 0 | 0 |  |  |
|  | - | - | - | - | Vaginal | 39.091 | Bifidobacterium vaginale | -8.95 | 2600 | 1537 | 648 | 0 | 213 | 0 | 200 | 26 | 0 | 0 |  |  |
| Tacrine | *H. sapiens* | Acetylcholinesterase | 7XN1 | ACHE | - | - | - | -5.51 | 162 | 40 | 192 | 1 | 0 | 0 | 37 | 30 | 1 | 0 |  |  |
|  | - | - | - | - | Gut | 33.88 | Raoultella ornithinolytica | -5.3 | 130 | 36 | 148 | 0 | 0 | 0 | 32 | 27 | 1 | 0 |  |  |
| Empagliflozin | *H. sapiens* | Sodium/glucose cotransporter 2 | 7VSI | SCL5A2 | - | - | - | -10.78 | 3178 | 1903 | 757 | 161 | 273 | 49 | 242 | 0 | 36 | 1 |  |  |
|  | - | - | - | - | Gut | 35.614 | Parabacteroides sp003473295 | -5.58 | 160 | 193 | 36 | 155 | 42 | 49 | 36 | 0 | 36 | 1 |  |  |

**Figure 4.**
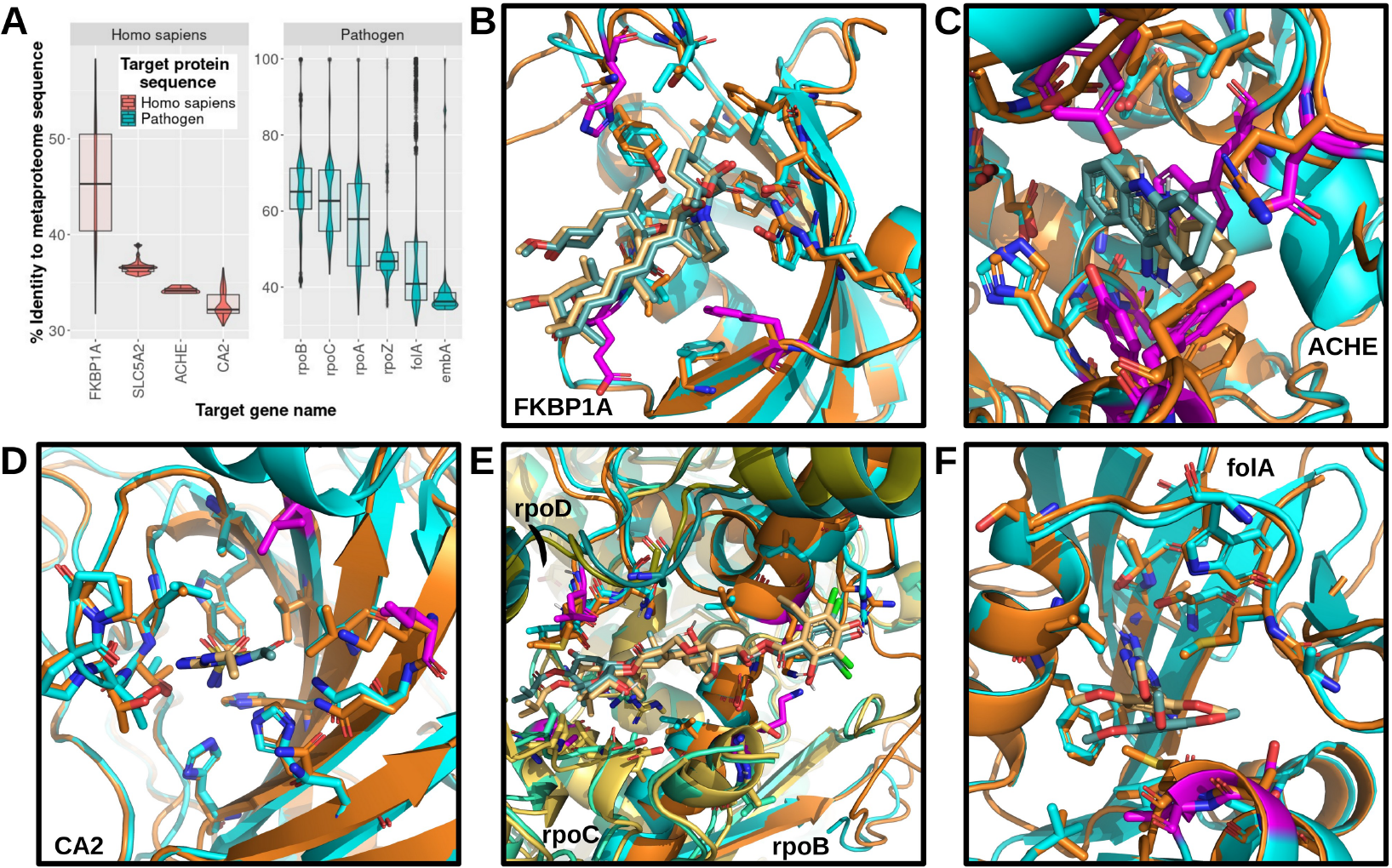
Protein-drug interactions The percent sequence identity distributions of microbial sequences aligned to drug targets with structural information related to drug binding are shown as a combined violin and box plot (A). The plots are colored and split into facets by target classification. The difference between protein-drug interactions in drug target (cyan) and microbiome proteins (orange) are depicted (B-F). Differences in aligned residues are colored in magenta on the drug target protein structure. The interacting residues between rapamycin and FKBP1A (B), tacrine and ACHE (C), acetazolamide and CA2 (D), fidaxomicin and DNA-directed RNA polymerase subunits rpoB, rpoC, and rpoD (E), and trimethoprim and folA (F) are shown as sticks.

#### 2.2.1. Human targets

In descending order of average sequence identity to the metaproteome sequences, the human targets consisted of peptidyl-prolyl cis-trans isomerase FKBP1A, sodium/glucose cotransporter 2 (SCL5A2), acetylcholinesterase (ACHE), and carbonic anhydrase 2 (CA2). Inspection of the interactions between rapamycin and the FKBP1A models revealed three single amino acid differences in the selected microbiome protein drug binding pocket (Figure 4B). Notably, these differences correspond to one residue insertion and to nonsynonymous variants, which may partially explain the decrease in predicted affinity of the microbiome FKBP1A to rapamycin compared to the human FKBP1A (Table 1). Interestingly, although SCL5A2 was second highest among the four human targets in average sequence identity, there were numerous variations in binding pocket residues, and the predicted affinity of empagliflozin with the microbiome SCL5A2 was approximately half of that with the human SCL5A2. The virtual docking of tacrine to the selected microbiome protein similar in sequence to human ACHE revealed a notable difference in the binding pocket: a tryptophan in the human ACHE exhibits pi stacking with tacrine, while this tryptophan is missing in the microbiome protein (Figure 4C). Of note, this and all other microbiome proteins that mapped to the human ACHE were predicted to be para-nitrobenzyl esterases. The non-overlapping putative function likely explains the differences in active site residues and, potentially, drug binding. Although the microbiome CA2 sequences were the lowest in sequence identity to the human CA2, the predicted affinities were almost identical and only 2 out of 15 binding site residues were different (Figure 4D). Remarkably, the human targets that mapped to the metaproteome sequences with the highest and lowest sequence similarities were found to have the most conserved binding pocket residues, while the binding pockets of the middle two were less likely to share structural features that enable high affinity interactions with the same drugs.

#### 2.2.2. Pathogen targets

In decreasing order of average sequence identity to the metaproteome sequences, the pathogen drug targets included DNA-directed RNA polymerase subunits (rpoA, rpoB, rpoC, rpoZ), dihydrofolate reductase (folA), and Probable arabinosyltransferase A (embA). The available structure of the target DNA-directed RNA polymerase (PDB: 7l7b) shows that fidaxomicin is bound to three subunits (rpoB, rpoC, and rpoD). Therefore, the protein assembly was modelled using the sequences from the same MAG. Among the three subunits, a comparable predicted affinity and only four total single residue variants were discovered between the target (*Clostridium difficile*) DNA-directed RNA polymerase and that of the human microbiome (Figure 4E). Comparing the binding pockets of the microbiome and target (*Mycobacterium tuberculosis*) embA revealed only one single amino acid difference, although several residues surrounding and leading into the binding pocket were not identical. Two binding pocket residues were different between the target (*Escherichia coli*) and microbiome folA proteins (Figure 4F). The predicted binding affinities for all three pathogen targets were comparable to their modelled microbiome counterparts. Overall, the pathogen targets were found to contain less structural and functional changes in the binding pockets between species even at approximately 30% global sequence identity. These data support the notion that drugs targeting pathogens may bind more promiscuously than human targets to microbiota proteins.

### 2.3. Clinical implications of drug promiscuity in the human microbiome

Nearly every aspect of human physiology (e.g. weight management, cardiovascular health, gut-brain axis) has been linked to the maintenance of microbiome composition and functional capacity[38]. The off-target effects of both antibiotics and non-antibiotic drugs on the human microbiome has been linked to various and diverse clinical manifestations[39]. Therefore, the data above may be used to better understand the effect of non-selective drug targeting on 1) the non-pathogenic bacteria in the gut, oral, and vaginal microbiomes and 2) the presence side effects when comparing drugs that do or not target proteins similar to the microbiome metaproteomes. Bartlett *et al*. [40] curated a list of 1,513 bacterial species that have been recorded to lead to infection, even considering opportunistic infections. Additionally, the SIDER Side Effect Resource [41] presents all recorded side effects, as defined in the MedDRA [42], associated with the use of thousands of drugs. A better understanding of the clinical implications may help us better untangle the complexity of pharmacological promiscuity and guide clinical decision making for personalized cases based on microbiome data.

#### 2.3.1. Non-selectively targeted pathogenic and non-pathogenic bacteria

Among the pathogenic species listed by Bartlett *et al*., 285 and 313 were found to be non-selectively targeted by human and pathogen drug targets, respectively (Figure 5A). A total of 1,876 and 1,987 other bacterial species – referred to as non-pathogenic bacteria – were mapped to the human and pathogen drug targets, respectively. Interestingly, the number of affected species between the human and pathogen drug targets were similar across the gut, oral, and vaginal microbiomes. Notably, more than half of the affected species are non-pathogenic, although the proportions are different for the gut microbiome than the oral and vaginal. The disproportionate number of non-pathogenic species affected in the gut microbiome provides additional data that suggest that the gut microbiome may be more susceptible to non-selective targeting than the oral and vaginal microbiomes. The number of shared affected non-pathogenic bacteria between the microbiomes show a similar trend to the overall data, wherein the gut and oral microbiomes have more overlapping off-target species (Figure 5B). In summary, diverse and nearly equal numbers of total non-pathogenic species that are found to be potentially non-selectively targeted were found for both human and pathogen drug targets. These data draw attention to the necessity for target specificity in drug design, discovery, and development.

**Figure 5.**
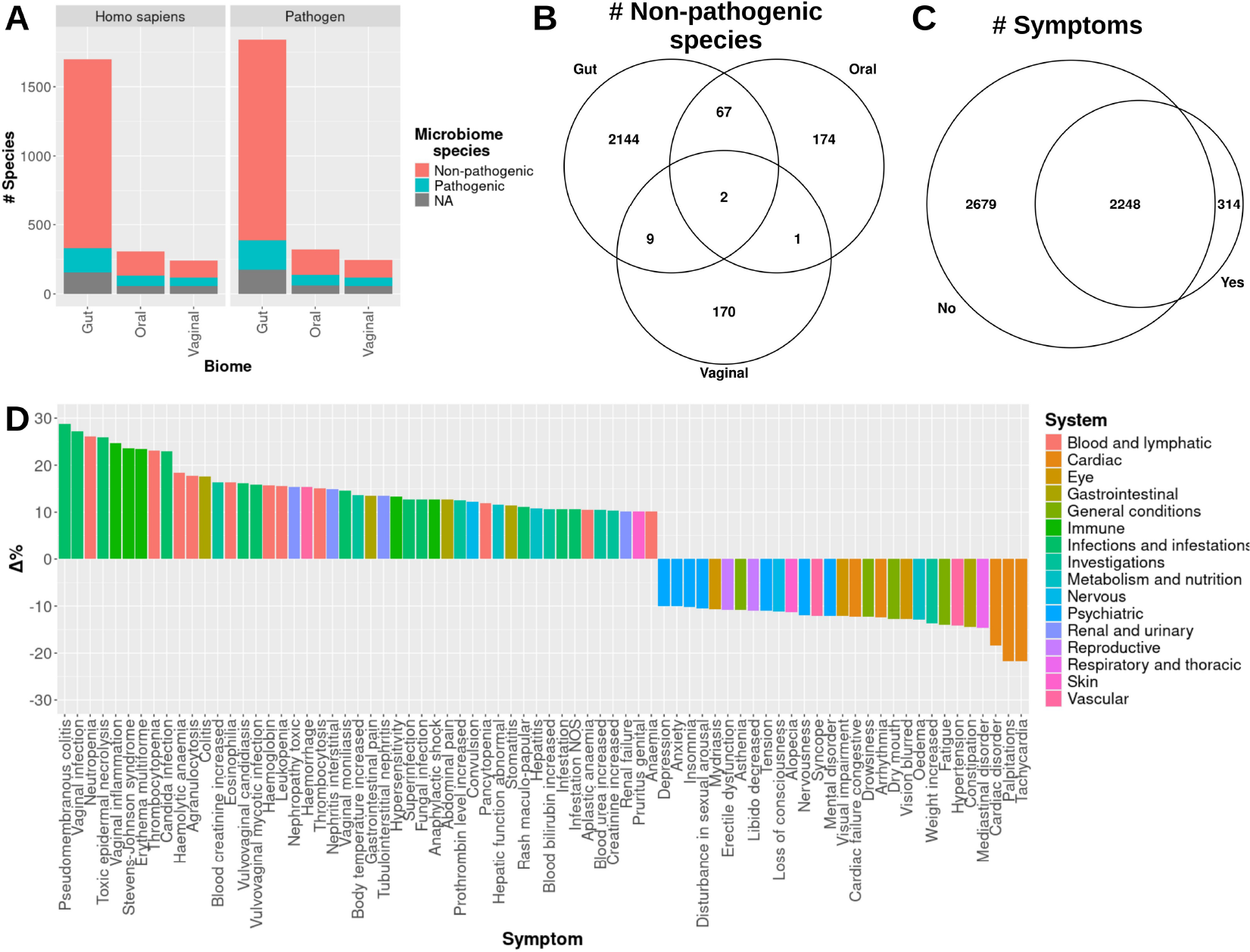
Clinical associations with drug target-metaproteome sequency identity A bar plot is presented that shows the number of pathogenic, non-pathogenic, and unclassified (“NA”) species that are similar to drug target sequences (A). The columns are split to represent the number of species per human and pathogen drug target per metaproteome. A Venn Diagram of the number of shared non-pathogenic species non-selectively targeted by each microbiome is shown (B). A Venn Diagram of the shared symptoms related to drugs that were found to be similar (“Yes”) and not similar (“No”) to microbiome metaproteomes (C). The percent changes in normalized prevalence of symptoms shared between “Yes” and “No” are plotted and colored by affected high-level organ system or disorder classification (D). Only symptoms with more than a 10% change are shown.

#### 2.3.2. Differential side effects of drugs based on non-selective of targeting microbiome proteins

The 126 and 611 drug targets that did and did not map to the microbiome metaproteome sequences were assigned as microbiome-affecting and non-microbiome-affecting, respectively. Upon filtering the SIDER datasets for side effects corresponding to the drug and drug target, 314 symptoms were unique to the targets that mapped to microbiome proteins, while 2,248 were shared (Figure 5C). Among the unique microbiome-affecting symptoms, 42 (13%) were related directly to infection and inflammation, such as fungal and bacterial infections, while the others were distributed among the remaining 25 high-level MedDRA System classifications. Notably, the symptoms of wound complications and surgical site reaction – potentially affecting the skin microbiome – were also found exclusively among drugs that non-selectively target microbiome proteins. Additionally, an analysis of the difference in prevalence among the shared symptoms may also reveal the specific effects of off-target effects on the microbiome. When comparing the percentage of prevalence of shared symptoms among drug targets that are and are not similar to the metaproteome sequences, certain MedDRA Systems seem to be particularly enriched (Figure 5D). Interestingly, several Infection (12 / 48), Blood and lymphatic (11), and Immune (5) symptoms were at least 10% more prevalent in microbiome-affecting targets. The prevalence of Cardiac (5) and Psychiatric (7) symptoms was particularly enriched for drugs that do not non-selectively target microbiome proteins. These data suggest that specific symptoms or organ system conditions may be more affected by drugs that non-selectively target the human microbiome.

## 3. Discussion

The effects of numerous drugs on the human microbiome have been widely shown in various studies[22,43–47]. Considering that different surfaces on the human body are inhabited by phylogenetically diverse microbes, there may be structural and functional overlap between current drug targets and microbial proteins. However, to date, no study has assessed the similarity between established drug targets and the proteins expressed by bacteria in different human microbiomes. Therefore, herein, sequence and structure bioinformatics methodologies were utilized to investigate how sequence identity between drug targets and microbiome proteins relates to structural features and, thus, drug promiscuity. These data were then combined with symptoms related to drugs and drug targets to gain insight into potential clinical outcomes associated with non-selective targeting of microbiome bacteria.

The average sequence identity between pathogen drug targets and the metaproteome of the gut microbiome, in particular, was notably higher than that of the drug target and microbiome associations. The significantly higher diversity of gut microbes compared to other human microbiomes has been noted previously [20], which, in combination with the findings in this study, suggest that a higher diversity may necessarily increase the likelihood that off-target drug targeting may occur. Furthermore, a higher number of non-selectively targeted species were found in common between the gut and oral microbiomes, which is not surprising considering their physical connection. The overarching differences in composition of off-targeted bacterial phyla between the gut/oral and vaginal microbiome metaproteomes highlights the diversity of species potentially targeted by drugs despite the target organism. The widely affected targets warrant further investigation into how drug targets for similar proteins may be affecting microbiome health.

These data suggest that a sequence identity of 30% may be used to determine similarity between drug targets of diverse phylogenetic ancestry. Notably, the pathogen target sequences were found to be more correctly mapped to microbiome protein sequences with identical functional annotations than the mappings of human target sequences. All three pathogen targets that were investigated using protein and drug structure analyses revealed consistent features to microbiome proteins, whereas the human drug targets were more dissimilar. Importantly, however, the human drug target with the lowest sequence identity (of the four with structural information) to the microbiome protein reported the most stable protein-ligand interaction compared to the other human drug targets. Incorporating structural information into drug target similarity is, therefore, crucial to accurately assess how drugs may interact with proteins that are homologous to the target. Variations in residues outside of the binding site may also affect binding site dynamics, and thus may guide drug design efforts. Of note, several hypothetical proteins were found to be similar to the drug targets, which may add putative functional information for these genes. The sequence, structural, and functional similarities between established drug targets and proteins in the human microbiome, as exhibited in this study, suggest that such checks should be routinely considered during target selection for drug discovery campaigns.

These results support previous findings that the microbes from various human micro-environments may be influenced by drugs taken via different routes of entry and, thus, have widespread effects on host-microbiome interactions[48]. An increase in the prevalence of side effects classified under immune system and blood disorders and infections were specifically linked to drugs that were found to non-selectively target microbiome species. These findings are interesting since drug-induced perturbations of the human microbiome have been shown to result in the proliferation of opportunistic bacterial pathogens[49,50]. Microbiome diversity forms the basis for maintaining protection against colonization by opportunistic pathogens [51]. Therefore, promiscuous drugs may disrupt the healthy microbiome diversity – as evidenced by the diverse number of phylogenetically-distinct non-pathogenic microbiome species with similar protein sequences to both human and pathogen drug targets – and give rise to micro-environments suitable for opportunistic pathogens[52]. A drug-induced expansion of opportunistic pathogens would support a recent report stating that bacteria in the patient microbiome are common causative agents in surgical site infections, as also seen in the side effect associations explored in this study[53].

Importantly, the reported associated side effects could be a result of other numerous and complex interactions that occur either solely in the human body or, perhaps, partly in connection with microbiota[54]. Considering that the potential off-target effects analyzed in this study were only assessed in the context of similar drug targets and not the vast protein space that may encompass the true landscape of drug promiscuity, further investigation is required to determine the side effects that are uniquely or differentially prevalent for certain drugs and drug targets[55]. Furthermore, drugs may be altered through biotransformation in microbial communities and, thus, affect downstream processes unrelated to the original target protein[56]. Bacteria non-selectively targeted may also be under evolutionary pressure to develop genetic mutations that nullify the effects of the drug, which may be transmitted horizontally to other microbiome bacteria and, thus, promote the spread of antimicrobial resistance mechanisms[57]. The impact of diet on both the microbiota and their combined effect on drug metabolism and off-target binding are also to be considered in the analysis of side effects[58]. Nevertheless, the preliminary clinical associations from this study suggest that certain side effects may be more prevalent when non-pathogenic microbiota are targeted.

Several factors have been suggested to help guide the selection of both human and pathogen targets for drug discovery efforts. In addition to the commonly performed sequence homology check between targets and human protein sequences, the data presented in this study point to the relevance for including a protein sequence identity screening for unintended microbiome targets. Artificial intelligence and machine learning approaches may also help further design drugs that are selective for specific targets and minimize off-target microbiome and human effects[59,60]. Alternatively, such data may be leveraged to target specific bacteria to reverse disease states that may be caused by phylogenetically related or distinct pathogens[61,62]. Notably, many of these factors can be applied when considering genes or proteins for diagnostic or prognostic measures as well[63]. In summary, the non-selective targeting of microbiome proteins may be minimized by ensuring that the intended protein target sequence lacks identity to metaproteome sequences. Further assessment of drug promiscuity among non-pathogenic microbes may inform how sequence, structural, and functional information can be utilized to minimize off-target effects.

## 4. Methods

### Data Collection and Analysis

The drug target protein UniProt [64] accessions were obtained from Santos *et al*. [26] and the protein sequences from UniProt. Only UniProt “Reviewed” drug target accessions were selected. The Gut v2.0.2, Oral v1.0.1, and Vaginal v1.0 microbiome metaproteome sequences, predicted functions, and phylogenetic lineages were downloaded from the MGnify database[27,34]. BlastP v2.6.0+ was used to map the drug target sequences to the microbiome metaproteome sequences with an e-value cut-off of 10^-6^[35]. Drug names and drug chemical characteristics, e.g. pKa, were obtained from PubChem [65] and DrugBank[66]. Drug side effects were retrieved from the SIDER 4.1 Side Effect Resource [41,67], and symptoms and organ system classifications were derived from MedDRA[42]. Protein and ligand structures were obtained from PDB[37]. Plots were generated using R v4.3.3[68].

### Protein structure modelling and ligand docking

Template-based protein structure homology modelling was performed using MODELLER v10.5 [69] with molecular dynamics-level optimization and refinement[70,71]. MAFFT was used to align protein structure sequences[72]. Structural templates for the microbiome protein modelling are listed in the “PDB” column of Table 1. In the case of tacrine, a hydrogen was added to the secondary amine based on its pKa value of 9.85 [73] using the attach function in PyMol v2.5[74]. CB-Dock2 [75,76] and MODELLER were used to perform template-based ligand docking (using the PDB structures in Table 1 as templates) of the drugs to the microbiome proteins. The drugs were also re-docked to the target protein as a control for affinity predictions. Protein-drug affinities and interactions were calculated using PRODIGY[77]. PyMol was used to visualize protein and drug structures.

## 5. Acknowledgements

CAB was supported by Antibiotic Research UK (ANTSRG 01/2019-PHZJ/687).

## 6. Author Contributions

All authors conceived and planned the experiments. CAB and SN carried out the experiments. All authors contributed to the interpretation of the results. All authors provided critical feedback and helped shape the research, analysis, and manuscript writing.

